# An expanding bacterial colony forms a depletion zone with growing droplets

**DOI:** 10.1101/2020.06.03.132498

**Authors:** H. Ma, J. Bell, J.X. Tang

## Abstract

Many species of bacteria have developed means to spread on solid surfaces. This study focuses on the expansion of *Pseudomonas aeruginosa* on an agar gel surface. We report the occurrence and spread of a depletion zone, where the layer of bacteria on the agar becoming thinner. The depletion zone occurs within an expanded colony under conditions of minimal water evaporation. It is colocalized with a higher concentration of rhamnolipids, the biosurfactants that are produced by the bacteria and accumulate in the older region of the colony. With continued growth in bacterial population, dense droplets occur and coalesce in the depletion zone, displaying remarkable fluid dynamic behavior. Whereas expansion of a central depletion zone requires activities of live bacteria, new zones can be seeded by adding rhamnolipids. These depletion zones due to the added surfactants expand quickly, even on plates covered by bacteria that have been killed by ultraviolet light. We propose a model to account for the observed properties, taking into consideration bacterial growth and secretion, osmotic swelling, fluid volume expansion, cell-cell interaction, and interfacial fluid dynamics involving Marangoni flow.

**Significance:** Bacterial growth and pattern formation have strong bearing on their biological functions, such as their spread and accumulation, biofilm growth & its effects on infection and antibiotic resistance. The bacterial species of this study, *Pseudomonas aeruginosa*, is a human pathogen responsible for frequent infections in wounds, airways, and urinary tract, particularly when involving the use of catheters. The findings of this study and the mechanisms we propose offer new insights on the important behaviors of bacterial collective motility, pattern dynamics, and biofilm growth.

## Introduction

In addition to planktonic existence, many species of bacteria have acquired an ability to colonize and spread on moist surfaces. While expanding on agar gel, a permeable, nutrient containing, semisolid surface, neighboring bacteria often move collectively, in packs [1], rafts [2], or large swirls [3]. When not mediated by functional flagella, collective bacterial spreading is referred to as “rafting” [2] or “sliding” [4]. In cases when the collective bacterial motility is aided by flagellated propulsion, this mode of surface motility is commonly defined as swarming [2, 5]. A popular depiction of swarming shows bacteria lined side by side, each driven by multiple flagella [5]. However, certain strains of flagellated bacteria with deficient or no flagella can spread with similar speed to that of their swarming counterparts [6–8], suggesting that a broader mechanism other than flagellated motility, such as sliding [4, 5] or volumetric fluid expansion [9, 10], may be the root cause for colony expansion. The roles of active flagella may include facilitating collective motion or fluid flow [5], and arguably, pumping fluids out of the agar or overriding surface friction [11].

A growing number of studies have attempted to explain features of bacterial colony expansion and pattern dynamics based on physics concepts and principles [8–10, 12–17]. From a physics perspective, a bacterial colony/swarm is a complex fluid containing active particles that grow in number. Coordinated motion, such as neighboring cells moving in packs, might simply be the outcome of a large number of densely packed cells spreading on a surface, through volumetric expansion driven by osmotic swelling [9, 13, 15], rather than by precisely controlled cell body alignment or direction of propulsion. Recent studies have provided measurements of cell density profiles [1, 12, 18] and fluid flow [19, 20] within large bacterial swarms, leading to an analytical model to account for the findings on these physical properties [17]. Another recent model based on the roles of surface forces predicts different patterns that may emerge in spreading bacterial colonies [10, 16]. A physical picture is starting to emerge that the governing mechanism for the spread and pattern evolution of a bacterial colony or swarm is an interplay between growth and interfacial fluid dynamics, involving osmotic and thin film flows [8, 20], wetting [10, 16], contact line pinning [8], Marangoni flow [12], and evaporation [15, 21].

In the meantime, species-dependent features and patterns have been widely reported. Proteus mirabilis, in particular, forms spatially periodic structures as the edge of its colony expands circularly outwards [22]. *Salmonella* [11] and *E. coli* [1, 19] tend to expand on agar with a smooth front. *Paenibacillus dendritiformis* [7] form dendritic protrusions, which can develop into highly branched structures. *Bacillus subtilis* [17, 23] and *pseudomonas* [15, 24–27] have been shown to expand in either smooth fronts or dendritic patterns dependent on various chemical and physical factors. We and others have shown, for instance, that by simply varying the agar concentrations [15, 26], adding external surfactants into the agar [15], or simply leaving the agar surface to dry for some time [15], the patterns of the colonies or swarms can vary from smooth to dendritic edges. Given the vast variation among species, and among even the same species under different conditions of experiments, bacterial patterns and forms remain a subject of ongoing study. The broader research community has not reached consensus concerning the underlining mechanisms, including to what extent they are biological or physical in nature [14, 28, 29].

This study sets a focus on *Pseudomonas aeruginosa*, a human pathogen with a rich variety of motility modes. It can swim, driven by its flagellum with a rotary motor at its base [30, 31], and can also drag itself over a solid surface by attachment and contraction of its Type IV pili [32–34]. When reaching high density on an agar surface, the flagellated bacteria display robust swarming motility [15, 25, 35]. Since mutants of *P. aeruginosa* with neither flagellum nor pili are still able to spread on agar [6], however, the species also manifests sliding motility [4]. In sessile state, *P. aeruginosa* is well known to form biofilms [36–40], thereby acquiring some essential biological functions such as mechanical resilience [41, 42] and antibiotic resistance [37, 43–47]. *P. aeruginosa* secrete biosurfactants and other extracellular polymeric substance (EPS) to facilitate cell-cell interactions, which are important for its swarming motility [25, 35, 40, 44, 48–50], as well as biofilm formation [35, 47, 51–53]. The biofilms of *P. aeruginosa* play essential roles on persistent infections [37, 47], which is a general issue on microbial infections [28, 54, 55]. The importance of *P. aeruginosa* as a pathogen, its different forms and numerous modes of motility may account for the extensive study of *P. aeruginosa*, including this work.

Here, we report new features observed on the largescale colony expansion of *Pseudomonas aeruginosa* on agar surface. With experiments performed under better humidity control, an important factor that caught our attention in recent studies [21, 56], we observe occurrence and spread of a depletion zone at the center of a large colony of *P. aeruginosa*, followed by growth and coalescence of bacterial droplets that display fluid dynamic behavior. We explore mechanisms to account for these features and converge to a model involving cell-cell cohesion, as well as thin film fluid dynamics. Our findings and interpretation offer new insights on bacterial collective motility, pattern development, as well as early stage biofilm growth.

## Methods

### Bacterial growth

Pseudomonas aeruginosa PAO1 (wild type) and its pilus-less mutant PAO1 ΔPilA, generously provided by Dr. Keiko Tarquinio of Emory University Medical School, were stored at −80°C in frozen medium containing 25% glycerol as cryo-protectant. A tiny amount was taken by scraping a frozen stock to grow in Tryptic Soy Broth (TSB) solution overnight in a flask at 37°C. We adopted a recipe from the Xavier lab [50] to prepare 0.5% agar plates. Specifically, the gel mixture containing 48 mM Na_2_HPO_4_, 24 mM KH_2_PO_4_, 8 mM NaCl, 1 mM MgSO_4_, 0.1 mM CaCl_2_, 0.5% glucose, and 0.5% agar is autoclaved for 35 minutes, mixed with 5g/L casamino acids (Bacto), pre-cooled to about 55 °C, poured in 50 mL volumes to large petri dishes of 15 cm diameter and placed in a bio-safe cabinet with lids open for 30 minutes to form 0.5% agar gel at room temperature. Then, 2.5μL of bacterial growth in Tryptic Soy Broth (TSB) was deposited at the center of the agar plate. The inoculated plate was placed inside a custom-built incubator with transparent top cover. The incubator maintains constant temperature at 37°C with changes within 0.5°C, controlled by a feedback sensor. Another sensor with a moisture release and a 3W mini-fan controls the humidity within the incubator to 60%. A Canon 7D DSLR camera was mounted on a stand so that pictures can be taken above the incubator without altering the temperature and humidity within it.

### Fluorescence Imaging

We designed and assembled a light box with illumination by LED lights on the sides (detailed in Supporting Material) in order to detect the distribution of rhamnolipids secreted by *P. aeruginosa* on the large agar plate. We added into agar gel Nile red, a dye that becomes visible when bound to rhamnolipids [33, 57]. The design works as the dye molecules enter the swarm fluid due to osmotic flow that fuels the bacterial colony expansion. Nile red, purchased in powder form from Fisher Scientific, was dissolved in methanol to the concentration of 1mg/mL [33]. The dye solution was added into agar mix in 1:100 (by volume) and then poured to a petri dish (diameter 15 cm) to form agar containing the dye. This gel plate was inoculated with PAO1 (wild type) or its mutant strain, ΔPilA, and grown in a dark incubator at 37°C and 60% humidity. A set of photos was taken each hour using a colored camera with RGB readings.

To obtain a radial fluorescence distribution plot, we split the round petri dish area in the photo to 1000 concentric annuli and calculated the average value of fluorescence intensity in each ring. The numbers of annuli from 1 to 1000 is scaled to radius of the petri dish from 0 to 7 cm. Then, fluorescence intensity-radius curves were plotted for photos shown at 3 time points.

## Results

### A depletion zone occurs at inoculation site hours following colony expansion

Following spot inoculation of wildtype *Pseudomonas aeruginosa* PAO1, the region containing the bacteria becomes visible in several hours. By 12 hours, the colony starts expanding notably beyond the initial area (Figure 1, with corresponding time lapse Movie S1 linked to Supporting Material; a similar example is also shown there as Figure S1). By 20 hours, a region of depleted bacterial density appears from the center. In the next 10 hours, the depletion region spreads behind the colony edge with a few hours in lag time, until it covers the whole plate. While the depletion zone expands, bacteria aggregate along its boundary, increasing the thickness of the bacterial film there. Meanwhile, droplets are shed when the depletion zone expands. Those droplets reside inside the depletion zone, migrating dynamically on the agar surface and coalescing with each other. Finally, the plate is covered by a film of seemingly dormant bacteria, with only a few large droplets stuck on the plate surface. Another example is shown in Figure S1 in the Supporting Material, with similar timeline and features. One notable exception in the 2^nd^ sequence is that the droplets appears later, emerging from within the depletion zone.

**Figure 1.**
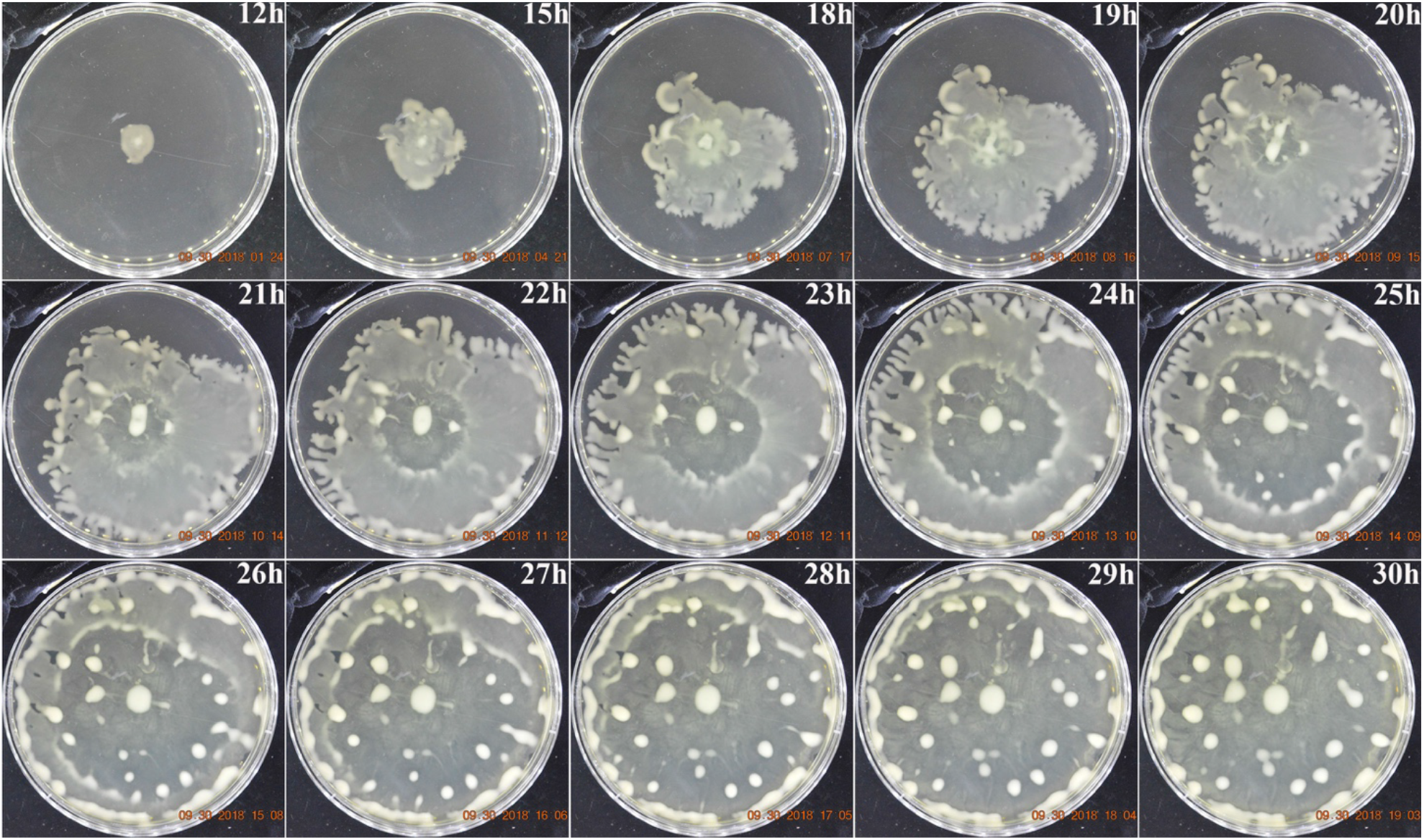
Expansion of *P. aeruginosa* colony over the course of 30 hours. The colony spreads over a large plate of 15 cm diameter in ~24 hours. A depletion zone occurs from the central region at ~20 hours and spreads over the entire plate in several hours. In the eantime, large droplets of bacteria form. They grow, migrate radially, and fuse with neighboring droplets. The growth took place on 0.5% agar on the covered plate, at 37°C, under 60% ambient humidity.

To test whether the depletion zone only occurs to the wildtype strain, we performed the same experiment using a mutant strain of PAO1, called ΔPilA, which expresses no pili. Our previous study confirmed that this pilus-less mutant spreads even better on agar than the wildtype [15]. Figure S2 in the Supporting Material shows a recorded sequence of the mutant colony’s spread, which features similarly prominent occurrence of a central depletion zone. We conclude that the depletion zone can occur on an expanding bacterial swarm independent of pili. In subsequent experiments, we focus on the wildtype strain in search of a common mechanism instead of looking for differences that are notable between different strains.

To test whether a depletion zone might always occur at the central area in a large colony irrespective of growth history, we inoculated an agar plate with the bacteria around its edge so that the population filled the plate surface from the edge inwards. We found that at about the same time as the colony reached the central region, roughly 18 hours into the colony spread, several depletion regions occurred near the outside edge (Figure S3), where the colony had been located several hours previously. This result indicates that indeed it is the growth history, not the exact geometry or shape of the colony that dictates where a depletion zone occurs. For convenience, all subsequent experiments were performed by point inoculation at the plate center.

### The shape of a depletion zone tracks that of a growing colony with several hours of delay

We noted that a depletion zone typically resembles the shape of a spreading colony. In Figure 2, for example, three pairs of dashed lines are drawn on corresponding pictures with consistent colors to show matching patterns between colony edges and depletion zones six hours later. This sequence of images shows the depletion zone tracking the colony edge with a constant time lag. In Figure 2f, the areas of the depletion zone and the colony are plotted versus time, showing essentially the same curve with a shift in time by about six hours. The traces of the depletion zone, viewed as the second wave of spreading, track that of the colony edge.

**Figure 2.**
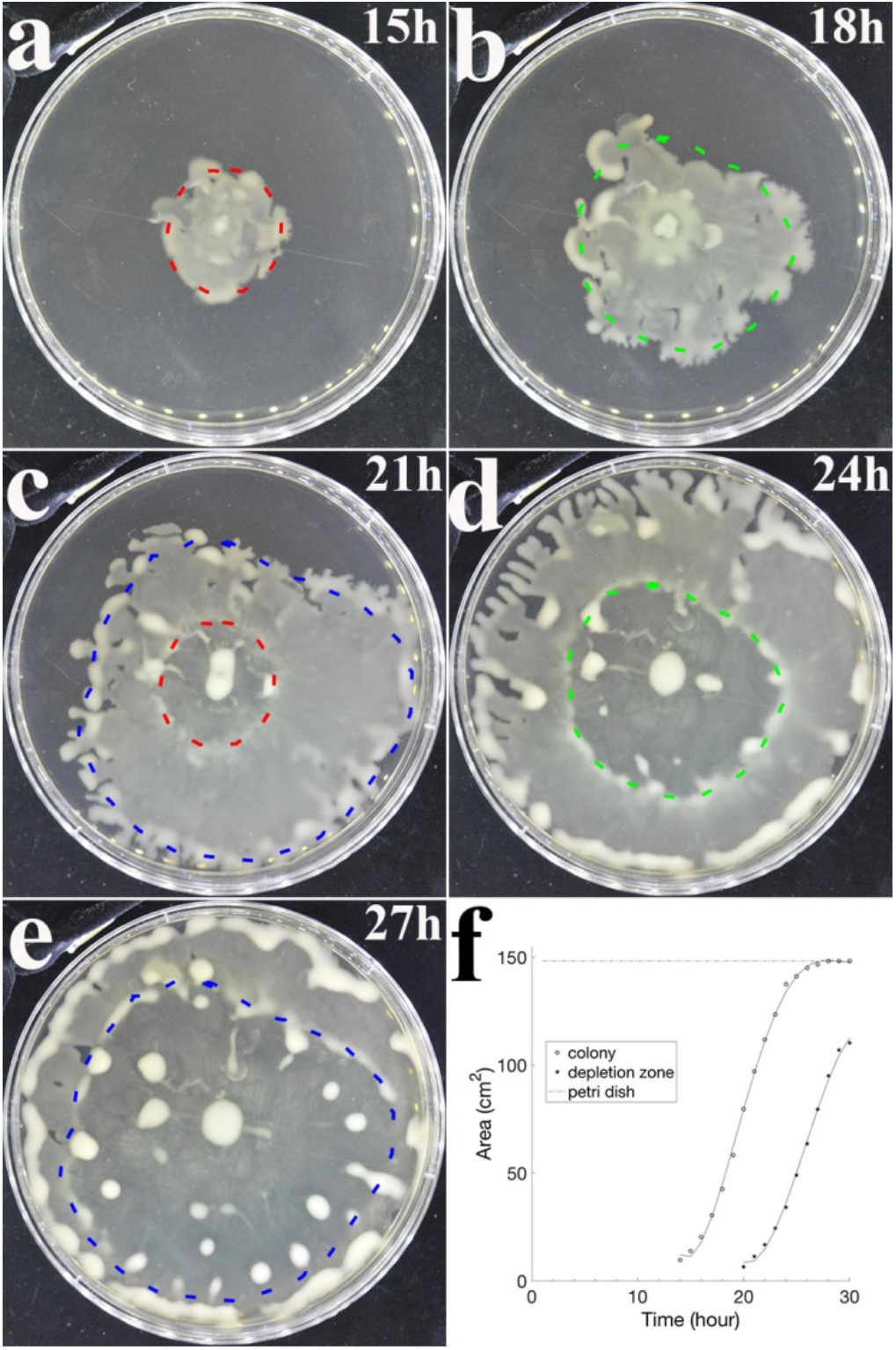
Occurrence and spread of a depletion zone at the central region of an expanding colony. The depletion zone, which was shown in Figure 1 to start about 20 hours after inoculation, tracked the colony covered area with about a 6-hour delay. Note the dotted contour lines in matching colors between vertically adjacent images. The tracking of the areas is most notable as they are plotted versus time in comparison at the bottom right. The plate diameter is 15 cm. The bacteria colony was inoculated on 0.5% agar gel, incubated at 37°C and under 60% humidity surrounding the covered plate.

In cases when a colony spreads into irregular shapes, and often with dendritic protrusions (such as one shown in Figure S1, for instance), the depletion zone resembles the rough contour of the colony several hours back, but is devoid of fine features. This is not surprising, since steep gradients of chemicals within a fluid layer are expected to be blunted by diffusion over several hours. We also performed a control experiment with no age variation within a large colony, by covering the plate surface with a thin layer of bacteria-containing medium. As expected, no depletion zone formed as the bacterial density increased uniformly over the entire plate (images not shown).

### The depletion zone tracks the region with higher concentration of rhamnolipids

We hypothesize that rhamnolipids, the bio-surfactants secreted by the bacteria in residence, may be a key material that promotes the occurrence of depletion zone. Thus, we use Nile red, a fluorescent dye with high affinity to rhamnolipids [57], to visualize its spatial distribution in the colony. The agar with Nile red dissolved in it supplies the colony with the dye, along with nutrients and water. After rhamnolipids bind Nile red, they fluorescently emit red light when excited with green light. Figure 3 shows good overlap of a depletion zone with the area of enhanced fluorescence, suggesting that rhamnolipids indeed track the depletion zone. The more time bacteria occupied a location, the more rhamnolipids accumulated. The history of spot inoculation gives rise to the highest concentration of rhamnolipids at the center, decreasing radially outward.

**Figure 3.**
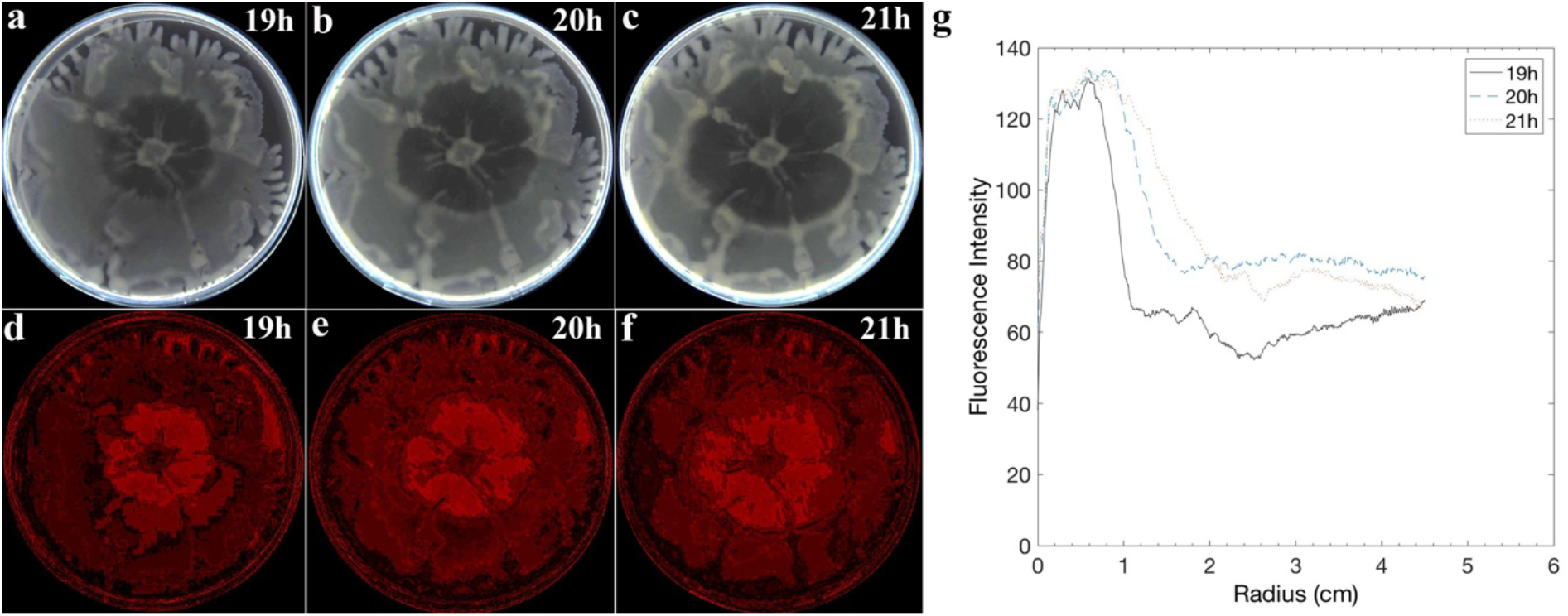
Imaging of rhamnolipid profile in a colony spread over an agar plate. **a.-c.** Photographic images of a large colony with an expanding depletion region. **d.-f.** Images of fluorescent emissions from Nile red illuminated by green LED light. The bright red region indicates high rhamnolipid expression. This area colocalizes well with the depletion region. **g.** profiles of radially averaged fluorescence intensity as functions of radius. The broadening peak tracks the growth of the depletion region over the 2 hour interval.

## Arrest of depletion zone by UV treatment

We performed another experiment to test whether the expansion of the depletion zone requires a live bacterial population or whether the zone might continue to expand, once formed, due to purely physical mechanisms. To do so, we placed a plate containing a developing depletion zone under UV illumination for 98 mins. It was compared with a control plate kept in the same incubator with the exception that its exposure to UV light was blocked by two layers of aluminum foil. The results show that UV exposure stopped expansion of the depletion zone (Figure 4), suggesting that continued expansion of the depletion zone requires live bacteria and/or their continued secretion of surface-active materials, such as rhamnolipids.

**Figure 4.**
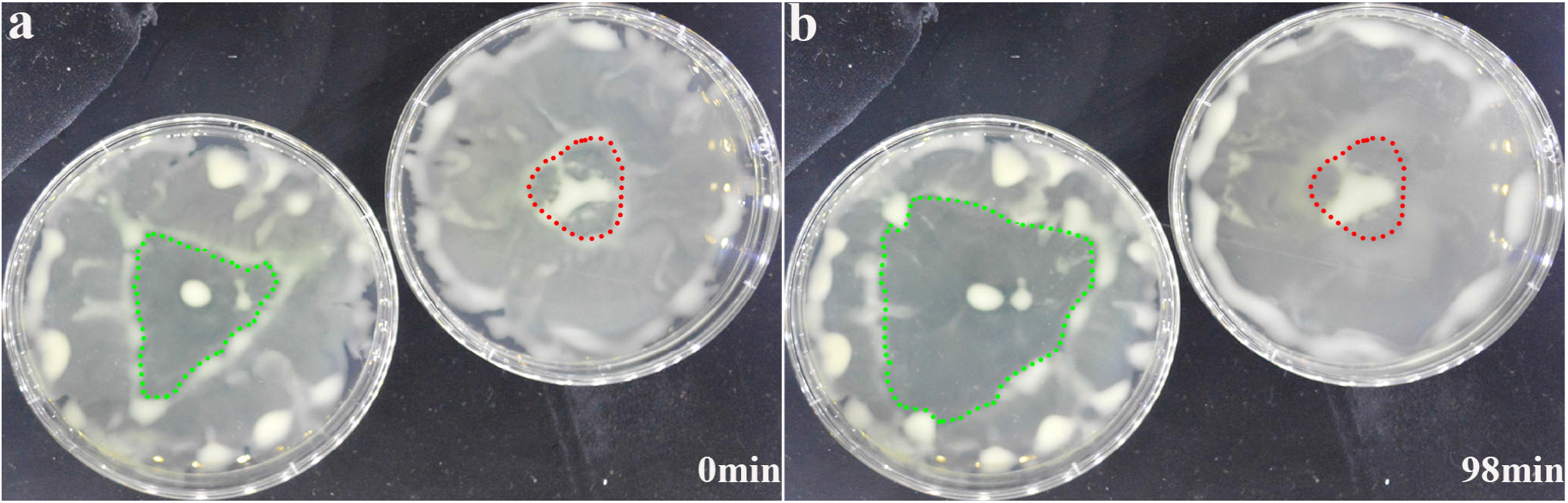
Arrest of a depletion zone by UV treatment. **a.** Depletion zones are noted on both plates fully covered by the bacteria grown over ~26 hours. **b.** The right plate was sterilized by UV light for 98 min while, as control, the left plate was blocked from UV damage. Both plates were kept in the same incubator. The dotted lines mark the boundaries of depletion zones formed in two plates. Note the depletion zone on the right plate stopped spreading due to UV sterilization, whereas on the control plate the zone continued to expand outward.

## Effect of surfactants on depletion zone

We performed an additional experiment to test whether added surfactants could induce a depletion zone. The effect of added drops of liquids containing 0.5% rhamnolipids were compared to that of water as control. This was done on both a live colony and one inactivated by UV illumination. The results show (Figure 5), surprisingly, that the added rhamnolipid drops germinate developing depletion zones under both conditions. In contrast, whereas water droplets containing no surfactant caused similar spots of notably diluted bacterial film initially, the spots did not expand over time. This result shows that a Marangoni flow due to surfactant density gradients on the colony surface is the likely cause of a depletion zone. In light of this finding, the observation shown in Figure 4 suggests that growth of a depletion zone depends on a high level of surfactants produced by live bacteria. Inactivated cells can no longer produce the high level of surfactants required to sustain the spread of the depletion zone. The observation of depletion zones geminated by surfactant droplets also rules out an alternative explanation to Figure 4 that loss of motility or cell death might be causing the bacteria to pack in a dense aggregate, thereby freezing the depletion zone.

**Figure 5.**
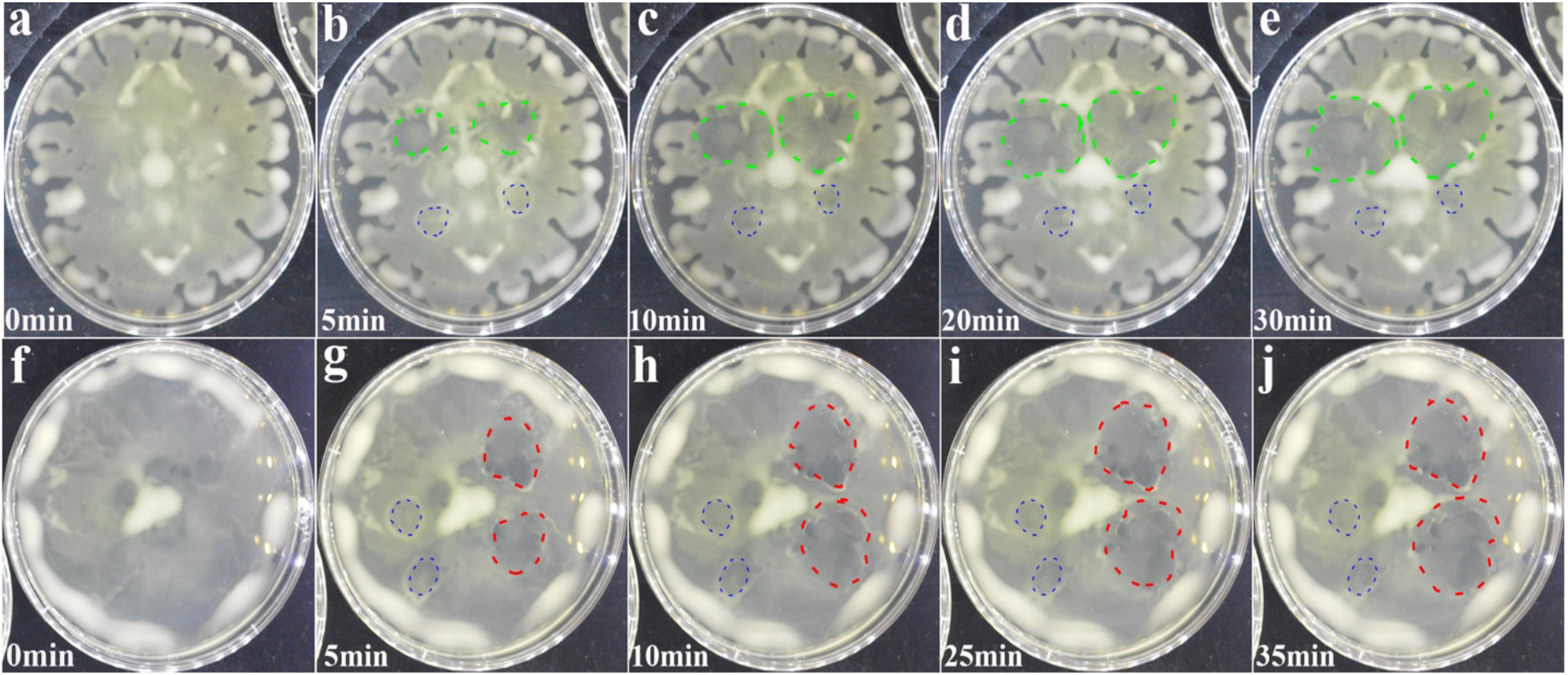
Comparison between plates covered by bacteria, either live or sterilized by UV after adding the liquid droplets. Bacteria are alive in the top row and sterilized in the bottom row. Two droplets each of water and rhamnolipids solution (10 μL) were added to the surface of each plate. Blue circles represent water droplets. Red and green circles represent rhamnolipids droplets (0.5%). Water droplets did not alter bacterial distribution in the area of addition, whereas rhamnolipids droplets induce small but expanding depletion zones on both plates.

## Discussion

Several physical conditions must be met for a bacterial colony to grow and spread over a nutrient rich agar gel. In order to take up nutrients from the agar for their growth, the bacteria must be able to secrete osmolytes and thereby draw fluids out of the agar. The rate of the volume increase must surpass loss due to evaporation in order to allow volumetric expansion of the bacteria-laden fluid. The bacteria must be able to produce surfactants in order to reduce the surface tension at the air-liquid interface and overcome the contact line pinning [8, 58, 59], so that the bacteria-containing fluid can readily spread over the agar surface. The fluid expansion may be further facilitated by Marangoni flow due to a surface tension gradient, which is caused by the higher concentration of rhamnolipids accumulated in the older region of the colony. Factors opposing fluid expansion on agar surfaces include contact line pinning [8, 58], a rapid rise of viscosity due to excretion of extracellular matrix in some species [8], and evaporation [21, 56]. Below, we discuss a few key observations made in this study in order to provide a model, which at least qualitatively accounts for the main findings in this report.

### Control of evaporation is a crucial factor in reproducible experiments on agar plates

Our recent experiments informed us that evaporation sensitively affects the bacterial colony expansion rate and the patterns formed [15, 21]. Although most microbiology experiments use covered agar plates and keep them in enclosed incubators or humidified chambers, for the sake of minimizing evaporation, we recognize through our recent studies that evaporation is always present while growing bacteria on agar plates. Most commercially available incubators are designed to allow enough air circulation to prevent condensation, which can easily form on the inside surface of a plate cover, for instance, due to even the slightest drop in temperature as soon as the plates are taken out for observation and/or imaging. Whereas the main function of the plate cover is suppressing evaporation by keeping the moisture inside, it is manufactured to have several bumps as spacers so that when it covers an agar plate, there remains a millimeter-thick gap of air for venting. Thus, the humidity inside the covered plate never approaches 100%. This means evaporation is a constant factor during incubation or observation. In this study, by keeping our incubation chamber humidified, typically at 60%, we found the rate of evaporation is significantly reduced as compared with placing plates in a commercial incubator. Note that the actual humidity inside the covered plate is much higher than 60% due to the large amount of agar inside, which contains over 97% water by volume. Since the venting only occurs around the thin gap on the edge, the actual humidity inside the large covered plate is also expected to have a gradient, reaching nearly 100% in the center, but gradually falling radially. The flanged plate cover ensures that the humidity inside is maintained at well over 90% even near the edge, which explains our observation that condensation frequently occurs on the inside cover if the outside humidity was increased to much above 60%, as soon as the temperature fluctuates by 0.5°C or more. Our limited testing suggests that better calibration and selection among various commercial incubators (including their stated operation settings, such as “gravity convection” versus “mechanical convection”, for instance, see https://assets.fishersci.com/TFS-Assets/LED/Specification-Sheets/PFCTMIDCTSN0214-MicrobiologicalIncubators.pdf) must be made in order for different microbiology labs to yield consistent results on bacterial colony growth and pattern observation. We conclude discussing this technical aspect by noting that effective reduction of evaporation using an incubating chamber built in house has led us to discovering new phenomena and gaining new insights.

### Fluidization and Marangoni flow jointly account for occurrence of a depletion zone

Our experimental findings show that a constant time lag exists between two spreading fronts, indicating the same spreading speed of the two fronts (Figure 2), and the region with high rhamnolipid concentration coincides with the depletion zone (Figure 3). The key property of constant time lag suggests that the spreading speed of the second front is not determined by dynamic parameters such as the local gradient of surface tension and viscosity. Instead, it is dictated by the altered rheological properties of the bacteria-laden film in a history dependent manner: after a fixed period of time since the local settlement of bacteria, the concentrations of rhamnolipids and possibly other extracellular matrix polymers reach threshold values. Some of the excreted molecules act as osmolytes, to pump fluid out of the agar in order to fuel the bacterial growth. The increased concentration of rhamnolipids, in the meantime, may facilitate the flow of the bacteria-laden liquid outward due to the well-known Marangoni effect [12], which predicts a flow along the concentration gradient of surfactants at the air-liquid interface. If the Marangoni flow is sufficiently strong, then, the majority of bacteria in the fluidized region might be carried by the flow and accumulate at the solid-fluid boundary [60]. Thus, the growth of the depletion zone may be limited by that of the fluidized region, instead of by the contact line spread at the outer edge. Further experiments are required to verify if the edge of the depletion indeed delineates a border between solid and fluid phases, as implied by the assessment above. For instance, a future study may be designed to directly measure locally the rheological properties of regions both inside and outside a depletion zone.

### An effective free energy model may predict droplet occurrence and growth

Large bacterial swarms have been previously modelled as a continuous thin-film fluid system [8, 10, 16, 18, 61, 62], instead of delving into the discrete, microscopic individuals. Extending the previous models, it may be possible to introduce an effective free energy as a framework in order to predict droplet formation based on minimizing the free energy. The proposed energy expression may include oxygen and nutrient density gradients over the thickness of the bacterial film [63, 64], which is estimated to be on the order of 100 μm based on previous studies [12, 57]. Thus, the model may include an explicit account for nutrient and/or oxygen transport within the bacteria-laden fluid film, as suggested previously [42, 47, 65]. Testing such a model would require measurements of nutrient and oxygen gradients over the thickness of the bacterial film, as well as factors secreted by the cells with spatial and time resolution, which is currently unavailable.

### A model of droplet formation based on cell-cell cohesion

Consistent with a free energy minimization scheme, one simple model for droplet formation may be developed that explicitly accounts for bacterium-bacterium cohesion. Specifically, these “free” bacteria may energetically prefer to gather and form densely packed droplets rather than to remain evenly dispersed in the fluid layer on the agar plate. This can occur due to an attractive cell-cell interaction, which could be mediated by extracellular matrix or cell surface polymers [51, 53], including surface layer proteins [65]. One can envision a transition from a thin film with cells uniformly dispersed to one with bacteria congregate into dense droplets. These bacterial droplets will grow at the expense of overall film thickness. Such a transition can occur when congregation of bacteria into large and dense droplets lowers the total energy of the system more than the energy cost of budding droplets out of the flat surface of a uniform bacterial laden film. This model implies that the bacterial droplets would only appear in the later period of the colony expansion, when the areal bacterial density reaches a high enough threshold. Since the model prediction varies with the cell-cell interaction strength and droplet surface tension, one could obtain an estimate of the cell-cell interaction based on measured bacterial film thickness when the droplets start to emerge, which sets an onset droplet size. For instance, the cell-cell interaction energy can be estimated as the surface energy of an onset droplet divided by the number of bacteria it contains. In essence, based on dynamic cohesion among motile bacteria, this simple model can account for the experimentally observed features. At the present, however, the values of relevant parameters, such as the surface tension, onset droplet size, and bacterial number within nascent droplets, are lacking, rendering the model qualitative in nature.

### A potential link between colony spread and biofilm growth

*P. aeruginosa* is known to form biofilms, which have been extensively characterized [36–40, 53, 57, 66]. At microscopic level, most of the published studies have shown images of the biofilms formed on glass or plastic surfaces, using mutant strains expressing green or cyan fluorescent proteins (GFP or CFP), which are amenable to confocal imaging (see, for instance, [38–40]). One study has shown that, intriguingly, biofilms formed by *P. aeruginosa* can detach upon aging or by raising the level of rhamnolipids, creating cavities on the scale of 100 μm [39].

The large depletion zone we observed, however, is different from the reported biofilm dispersal caused by rhamnolipids [39]. First, although the wildtype and ΔPilA strains are known to form biofilms, the time scale for that process is several days, much longer than the period for the features to occur in our study. Second, the central hollowing effect in aged biofilms, as reported by Boles and co-workers, occurs locally at numerous locations and on the length scale of 100 μm. In contrast, the depletion zone we observed occur to the entire colony, typically several centimeters in size. Last but not least, unlike in the biofilm growth experiment, the depletion zone occurrence we observed does not depend on adhesion of the bacteria to a solid surface. Because our experiments were performed on a large agar plate not amenable to microscopic imaging, however, we do not know in our study if cells actually adhere to the agar surface. The bacterial film in the depletion zone may still be rather thick based on the observation that bacteria-laden droplets continue to appear and grow in the region over time. Notably, our observations are on much larger sizes and thickness than the previous studies performed by microscopic imaging. In the future, we hope to be able to measure the thickness profile of the large bacterial colony spread over the agar plate, perhaps by adopting a method similar to that reported in [67].

## Conclusion

Bacterial growth and pattern formation have strong bearing on their biological functions, such as their spread and accumulation, biofilm growth [41, 47, 68] and its roles on infection [28, 54, 55], particularly, antibiotic resistance [37, 43, 45]. We report on observation of a depletion zone in an expanding colony of *Pseudomonas aeruginosa*, as well as occurrence and growth of bacteria-laden droplets. Our observation is explained by assessing bacterial growth, osmotically driven volume expansion, cell-cell cohesion, as well as interfacial fluid dynamics involving Marangoni flow. This report opens the door to more experiments as well as more comprehensive modeling in future work, taking into consideration all the essential parameters. By recognizing the robust interfacial fluid dynamics microbes must follow, new applications may be developed towards controlling, facilitating or inhibiting, when needed, the spread of microbes on various fluid-solid interfaces, which are highly relevant to biofilm growth and infection control. In addition, the new findings in this study suggest that better control of evaporation is required for consistency among common microbiology experiments using agar plates.

## Acknowledgement

We thank Dr. Keiko Tarquinio of Emory University Medical School, formerly of Brown University Medical School, for the bacteria stains. We thank Mr. Enrui Zhang and Prof. Huajian Gao for discussions.

## Author Contribution

J.X.T., J.B. & H.M. designed the study. H.M. & J.B. performed the experiments and analyzed the images and results. J.X.T & H.M. wrote the paper.

